# Infection with the chytrid fungus (*Batrachochytrium dendrobatidis*) may distort sex ratio via sex reversal but not sex-dependent mortality in a common Eurasian amphibian

**DOI:** 10.1101/2023.08.25.554788

**Authors:** János Ujszegi, Nikolett Ujhegyi, Emese Balogh, Zsanett Mikó, Andrea Kásler, Attila Hettyey, Veronika Bókony

## Abstract

One of the major factors driving the currently ongoing biodiversity crisis is the anthropogenic spread of infectious diseases. Diseases can have conspicuous consequences, such as mass mortality events, but may also exert covert but similarly severe effects, such as sex ratio distortion via sex-biased mortality or sex reversal. Chytridiomycosis, caused by the fungal pathogen *Batrachochytrium dendrobatidis* (Bd) is among the most important threats to amphibian biodiversity. Yet, whether Bd infection can skew sex ratios in amphibians is currently unknown, although such a hidden effect may cause the already dwindling amphibian populations to collapse. To investigate this possibility, we collected common toad (*Bufo bufo*) tadpoles from a natural habitat in Hungary, and continuously treated them until metamorphosis with sterile Bd culture medium (control), or a liquid culture of a Hungarian or a Spanish Bd isolate. Three months after metamorphosis we dissected the individuals and diagnosed their phenotypic and genetic sex. Bd prevalence was high in animals that died during the experiment but was almost zero at sexing. Survival was generally low in the control group, but it was further lowered by the Bd treatments. We did not observe sex-dependent mortality in either treatment. However, treatment with the Spanish Bd isolate significantly increased the frequency of sex reversal: 3 out of 9 genetic females developed into phenotypic males. Based on our results, Bd infection may have the potential to affect sex ratio in common toads through female-to-male sex reversal, but future research is needed to ascertain the generality of these findings.

## Introduction

The sixth mass extinction of Earth’s history is happening in our present days (Ceballos et al. 2015). One of the major factors responsible for this biodiversity crisis is the spread of infectious diseases in wildlife populations, both directly and indirectly facilitated by anthropogenic activities and human-induced environmental changes (Smith et al. 2009; Lindahl & Grace 2015). Though amphibians were not significantly affected in earlier extinction events, in the last century they became the most vulnerable taxon among vertebrates: more than 40% of amphibian species are threatened by extinction (Monastersky 2014; IUCN 2022). The globally spreading infectious diseases are among the main drivers of amphibian declines (Harvell et al. 2002; Pounds et al. 2006). Chytridiomycosis caused by the chytrid fungi *Batrachochytrium dendrobatidis* (Bd) and *Batrachochytrium salamandrivorans* (Bsal) (Van Rooij et al. 2015) has already led to the decline or extinction of several hundred amphibian species, due to repeated introductions arising from human activities (Lips 2016; O’Hanlon et al. 2018; Scheele et al. 2019). While the known geographic distribution of Bsal is restricted to relatively small areas (Spitzen-van der Sluijs et al. 2016), Bd is present on all continents apart from Antarctica. The waterborne motile zoospores of Bd infect only the keratinous epidermal layers of the amphibian skin (Berger et al. 1998). In case of serious infection in metamorphosed anurans, the structural damage caused in the skin can impair its osmoregulatory function leading to shifts in blood electrolyte balance. This may ultimately result in cardiac asystolic death (Voyles et al. 2009). Sublethal Bd infection can negatively affect life-history traits such as body mass, growth and development (Parris & Cornelius 2004; Blaustein et al. 2005; Garner et al. 2009; Hanlon et al. 2015), it may reduce the amount of skin-secreted chemical defences (Ujszegi et al. 2021) and often causes elevated levels of glucocorticoid “stress hormones” (Gabor et al. 2013, 2015). These sublethal effects can also contribute to increased mortality when combined with other stressors (Rohr et al. 2013; Mccoy & Peralta 2018).

When one sex is more sensitive to a stressor than the other one, sex-biased mortality can distort the sex ratio at the population level (Székely et al. 2014). Parasites and pathogens often have greater impact on survival in one sex than in the other (Moore & Wilson 2002; Moore 2003). When infectious diseases distort the sex ratio, this can exacerbate their negative effects on population viability (Rosa et al. 2019). Skewed sex ratios of breeding populations constrain effective population sizes and adaptive potential, and can have cascading effects on other species and even ecosystems (Mitchell & Janzen 2010; Edmands 2021), although they may also facilitate the evolution of sex-specific life histories and social systems (Schacht et al. 2022). However, whether Bd causes differential mortality in males and females is poorly understood, partly because identifying sex is not possible in most amphibians until they reach sexual maturity (Ujhegyi & Bókony 2020), except for a few species where molecular sexing methods are available (reviewed by Nemesházi & Bókony 2022).

Besides causing sex-biased mortality, stressors may also alter sex ratio by interacting with environmentally sensitive sex determination. In amphibians, phenotypic sex is determined by the sexual genotype (e.g. sex chromosomes) but can also be affected by environmental factors experienced during early ontogeny, causing sex reversal whereby individuals develop the phenotypic sex opposite to their genetic sex (Flament 2016; Nemesházi & Bókony 2022). Sex reversal may lead to skewed adult sex ratios reducing population viability (Stelkens & Wedekind 2010; Bókony et al. 2017; Nemesházi et al. 2021), although it may also facilitate adaptation to different environments (Geffroy & Douhard 2019). In ectothermic vertebrates, sex reversal can be triggered by endocrine disrupting chemicals and various stressful stimuli including low or high temperatures or pH, hypoxia, hyperosmotic conditions, crowding, starvation, and altered light conditions (Baroiller & D’Cotta 2001; Geffroy & Douhard 2019; Castañeda-Cortés & Fernandino 2021; Geffroy 2022). Although it has been shown in several invertebrates that parasites and pathogens can influence sexual differentiation (Rodgers-Gray et al. 2004; Narita & Kageyama 2008), whether pathogenic infection can induce sex reversal in vertebrates has never been studied to our knowledge.

In this study, we experimentally investigated the effects of larval Bd exposure on sex-biased mortality and on the incidence of sex reversal in common toads (*Bufo bufo*). This species is widespread and abundant across Eurasia (IUCN 2022). Toad tadpoles can be experimentally infected with Bd, and costs of Bd infection can reach measurable levels during early life stages (Garner et al. 2009, 2011; Woodhams et al. 2012; Ujszegi et al. 2021), while adults tolerate Bd presence well with low infection intensities (Baláž et al. 2014; Vörös et al. 2018). The larval phase is further relevant to our objectives because it is often in the early developmental stages that individuals are exposed to the waterborne Bd zoospores due to their aquatic lifestyle (Kilpatrick et al. 2010), and the larval phase also includes the period when sex is determined and gonad differentiation is sensitive to environmental perturbations (Nemesházi & Bókony 2022).

## Materials and methods

### Experimental procedures

In 2022, we made several attempts to collect common toad eggs from early spring on, but we always found only a limited number of eggs, often already frost-bitten. Therefore, we were forced to use wild-caught tadpoles for the experiment. We collected ca. 400 tadpoles in development stages 26-28 (Gosner 1960) in May using dip nets from a pond near Budapest, Hungary (Békás-tó: 47.57638° N, 18.86918° E). We transported the animals to the Experimental Station of the Plant Protection Institute in Julianna-major, Budapest, and initially held them in groups of 38-40 tadpoles in transparent plastic containers (27 × 18 × 14 cm) in 5 l reconstituted soft water (RSW; USEPA 2002). During the first week, we maintained 16.3 ± 0.3 °C (mean ± SD) temperature and set the lighting to match outdoor conditions. We fed the tadpoles with slightly boiled, chopped spinach twice a week. After one week, we haphazardly selected 300 healthy-looking tadpoles, and placed them in groups of ten into 5 l RSW using the same type of containers. We released the remaining tadpoles back into their original habitat.

From this point on, we exposed tadpoles during the entire larval development to sterile culture medium (control), or to liquid culture of one of two isolates of the global pandemic lineage (GPL) of Bd. The Spanish isolate (IA042) originated from a dead *Alytes obstetricans* collected in 2004 by J. Bosch (Biodiversity Research Institute, University of Oviedo, Spain) from a mass mortality event in Spanish Pyrenees. The Hungarian isolate (Hung_2014) was collected from a living *Bombina variegata* in 2014 by J. Vörös (Department of Zoology, Hungarian National History Museum, Budapest, Hungary) in the Bakony Mountains, Hungary. For details of Bd culturing and exposure see below. We used these two isolates because they had been available in our laboratory; we did not aim to compare the effects of the two isolates since their culturing history is very different. We exposed individuals in groups of ten to each of the three treatments in ten replicates, resulting in a total of 30 experimental units and 300 tadpoles. We randomly assigned containers to the treatments and arranged the 30 rearing boxes into 10 spatial blocks, each containing one replicate from each treatment. Temperature was 18.4 ± 1.5 °C (mean ± SD) during the experiment and we adjusted the lighting weekly to outdoor conditions. We fed tadpoles with spinach *ad libitum* and changed water twice a week using different dip nets for each treatment to prevent cross-contamination. Three weeks after starting the treatments, we replaced the rearing containers with larger ones (31 × 20 × 16 cm) and reared each group of 10 tadpoles in 10 l RSW from that point on to provide a larger water volume for growing larvae. Bd exposure lasted until the start of metamorphosis.

When an individual reached development stage 42 (forelimb emergence) we weighed it to the nearest mg and placed it into a covered transparent plastic container (31 × 20 × 16 cm). We filled these containers with 250 ml RSW and lifted one side by ca. 2 cm to provide animals with both water and a dry surface. Once the animals reached development stage 46 (total tail resorption), we placed them into transparent plastic containers (21 × 16 × 12 cm) covered with a perforated lid, lined with wet paper towels and a piece of cardboard egg-holder as a shelter. Each box contained 5 individuals from the same treatment group. After the completion of metamorphosis, we fed the toadlets *ad libitum* twice a week, initially with springtails (*Folsomia* sp.), then after three weeks with small crickets (*Acheta domestica*, instar stage 1–2) sprinkled with a 3:1 mixture of Reptiland (Art. Nr. 76280; Trixie Heimtierbedarf GmbH & Co. KG, Tarp, Germany) and Promotor 43 (Laboratorios Calier S.A., Barcelona, Spain) to provide vitamins, minerals, and amino acids. We recorded the dates of starting metamorphosis, completion of tail resorption, and eventual mortality. Dead specimens were conserved in 96 % ethanol. Ten weeks after starting the treatments, we euthanized tadpoles that had not started metamorphosis (N=32) in a water bath containing 6.6 g/l tricaine-methanesulfonate (MS-222) buffered to neutral pH with the same amount of Na2HPO4, and preserved them in 96 % ethanol.

Four months after starting the treatments, we euthanized all remaining juveniles in a water bath using the same method as for tadpoles. At that time, the toadlets were between 7 to 14 weeks old after metamorphosis; by this age the gonads of this species are fully differentiated and easy to observe (Ogielska & Kotusz 2004). We removed both feet of each euthanized individual and stored them in 96 % ethanol. We dissected the individuals, examined their internal organs under an APOMIC SHD200 digital microscope, and categorized phenotypic sex as male (testes) or female (ovaries). We did not observe any intersex individuals exhibiting both testes and ovaries. Dissected bodies with gonads were stored in 10 % formalin.

### Maintenance of Bd culture and exposure

Cultures were maintained in TGhL medium (mTGhL; 8 g tryptone, 2 g gelatine-hydrolysate and 4 g lactose dissolved in 1 l distilled water) in 25 cm^2^ cell culture flasks at 4 °C and passed every three months into sterile mTGhL. One week before use, we inoculated 110 ml mTGhL with 3 ml of these cultures in 175 cm^2^ cell culture flasks and incubated these for seven days at 21 °C. We assessed the concentration of intact zoospores using a Bürker chamber at ×400 magnification and diluted both cultures to an equal concentration. After each water change, we inoculated 10 ml of these cultures into the tadpole-rearing containers holding 10 l RSW, resulting in a mean initial concentration of ∼1940 zoospores/ml in the rearing water. We inoculated control containers with the same quantity of sterile mTGhL. Contaminated water and equipment were disinfected overnight with VirkonS before disposal (Johnson et al. 2003).

### Assessing Bd infection prevalence and intensity

We assessed infection prevalence and intensity from dissected mouthparts of preserved tadpoles and from toe clips of metamorphosing and metamorphosed individuals. The aim of these measurements was to infer if the Bd treatments were successful. To save costs, we randomly selected ca. one third of the individuals from the control group. We accidentally lost one sample from an individual euthanized as tadpole in the Hungarian Bd treatment group and several further samples of dead animals due to rapid decay (for analysed sample sizes per group, see Table 1). We homogenized tissue samples, extracted DNA using PrepMan Ultra Sample Preparation Reagent (Thermo Fisher Scientific, Waltham, Massachusetts, USA) according to previous recommendations (Boyle et al. 2004), and stored extracted DNA at −20 °C until further analysis. Following a standard amplification methodology targeting the ITS-1/5.8S rDNA region, we ran real-time quantitative polymerase chain reactions (qPCR) on a BioRad CFX96 Touch Real-Time PCR System (Bio-Rad Laboratories, Hercules, California, USA). To avoid PCR inhibition by ingredients of PrepMan, we diluted samples ten-fold with double-distilled water. We ran samples in duplicates, and in case of contradictory results, we repeated reactions in another duplicate. If it again returned an equivocal result, we considered that sample to be Bd positive (Kriger et al. 2006). Genomic equivalent (GE) values were estimated from standard curves based on five dilutions of a standard (1000, 100, 10, 1 and 0.1 zoospore genomic equivalents; provided by J. Bosch).

**Table 1:**
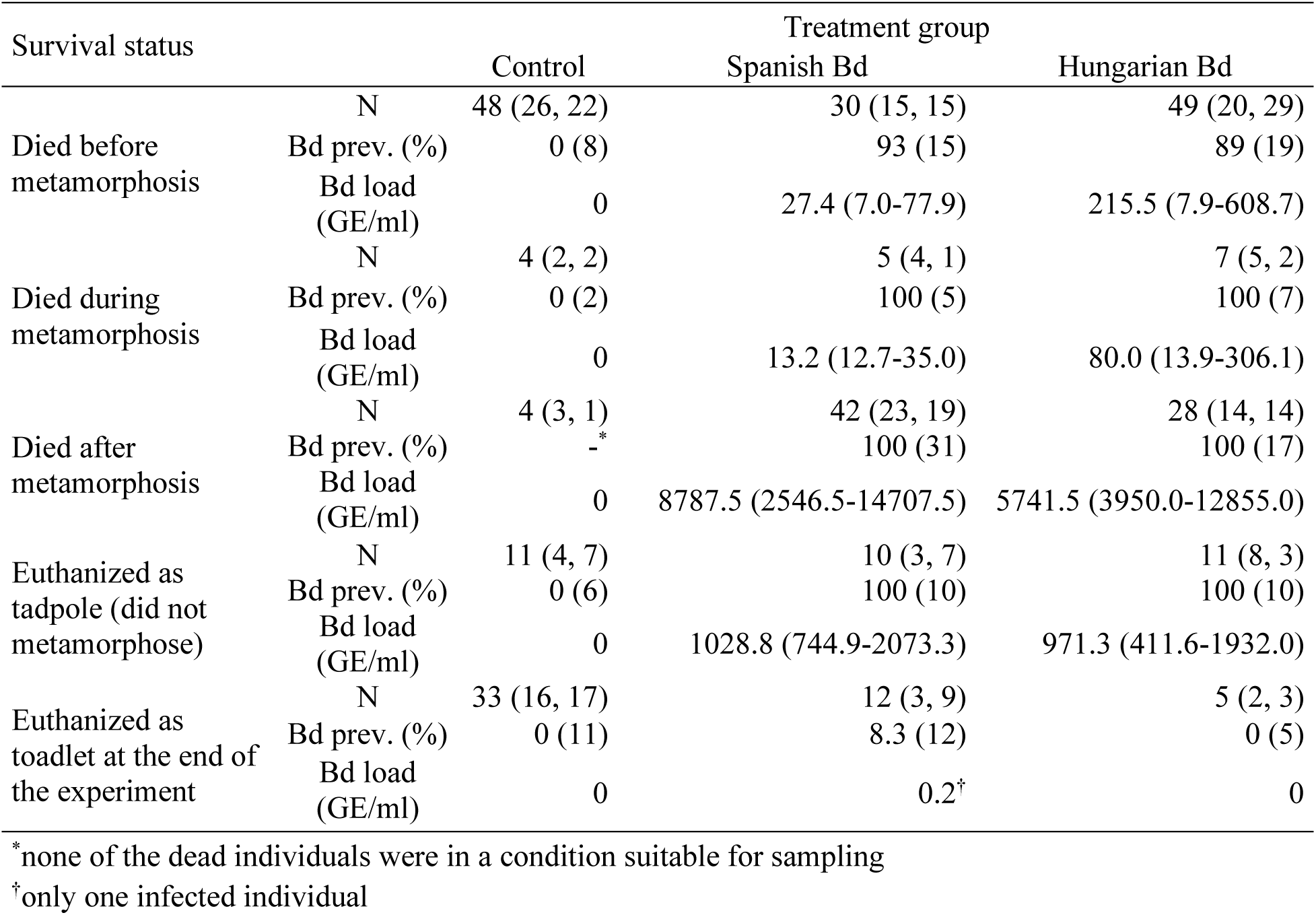
Survival data, Bd prevalence (Bd prev., with sample size of Bd-tested individuals in brackets) and Bd load (median, with interquartile range in brackets) in each treatment group. Sample sizes (N) of genetically sexed individuals are given as: total (male, female).

### Genetic sex determination

We extracted DNA from tissue samples using E.Z.N.A. Tissue DNA Kit following the manufacturer’s protocol (except that digestion lasted at least 3 hours). From animals that died prematurely as tadpoles or during metamorphosis, we used the tail, whereas from metamorphosed individuals we used one or two feet depending on size. For diagnosing genetic sex, we used a method published earlier (Nemesházi et al. 2022). For individuals that died before phenotypic sexing, we used two markers (c16 and c2). These markers yielded the same result for each sample. For phenotypically sexed individuals, we used marker c16, and when the result contradicted the phenotypic sex, we used two other markers (c2 and c5; both applied to two DNA samples isolated separately from each individual whenever it was possible). We judged an animal to have undergone sex reversal when all three sex markers contradicted phenotypic sex. In these latter cases, we confirmed phenotypic sex by histological examination of the formalin-stored gonads using routine histological procedures (Nemesházi et al. 2022).

### Statistical analyses

All analyses were run in ‘R’ (v4.0.5; R Core Team 2021). For the analysis of survival, we used Cox’s proportional hazards models (R package ‘survival’, function ‘coxph’; Therneau 2023). We treated all euthanized individuals (i.e. those that survived until the end of the experiment, and those that were euthanized because they did not start metamorphosis) as censored observations. We measured survival in units of life stage (i.e. before, during, and after metamorphosis) rather than in days, to maximize sample size (because the exact date of death was lost for a few individuals) and to accommodate time-dependent effects. Initial diagnostics suggested that the effect of Bd treatment was not constant over time, violating the proportional hazards assumption; therefore, we allowed for time-dependent treatment effects by stratifying the model by life stage. First (model 1), we included only the interaction between treatment and life stage, and we estimated the differences between the control and Bd-treated groups using the ‘emmeans’ function of the ‘emmeans’ package (Lenth 2018), correcting the P-values for false discovery rate (Benjamini & Yekutieli 2001). To test whether mortality was sex-dependent (model 2), we added the time-independent effect of genetic sex into the previous model. Then, to test whether the effect of treatment was sex-dependent (model 3), we added the two-way interaction between treatment and genetic sex and compared this model with model 2 using a likelihood ratio (LR) test.

To analyse female-to-male sex reversal, we used only genetic females, and we compared the proportion of phenotypic males of the two Bd treatment groups to the control group using Fisher’s exact test. We did not apply a more sophisticated analysis, nor did we analyse male-to-female sex reversal, because the overall sample size of toadlets surviving to phenotypic sexing was low, with zero individuals in certain combinations of treatment by sex. We did not analyse Bd prevalence and infection intensity statistically because the samples were taken whenever the individuals died; we used these data only to qualitatively assess if Bd treatment was successful.

## Results

None of the analysed control samples tested positive for Bd (Table 1). At the same time, Bd treatments were successful, as infection with both isolates resulted in high prevalences and infection intensities in individuals that died before the end of the experiment (Table 1). Out of the 50 toadlets that survived until the end of the experiment, only one had a detectable Bd load at dissection (Table 1).

Survival was low in the control group (Table 1), and it decreased further as a result of Bd treatment (Fig. 1). Specifically, mortality after metamorphosis was significantly higher in the groups treated with the Hungarian or the Spanish isolate compared to the control group (Table 2), whereas there was no significant treatment effect during the tadpole stage and metamorphosis (Table 2). Genetic sex did not affect survival either alone (model 2, female/male hazard ratio: 0.874 ± 0.119, P = 0.321; Table 1) or in interaction with Bd treatment (model 3, LR test: χ2 = 3.012, df = 2, P = 0.222; Table 1).

**Figure 1:**
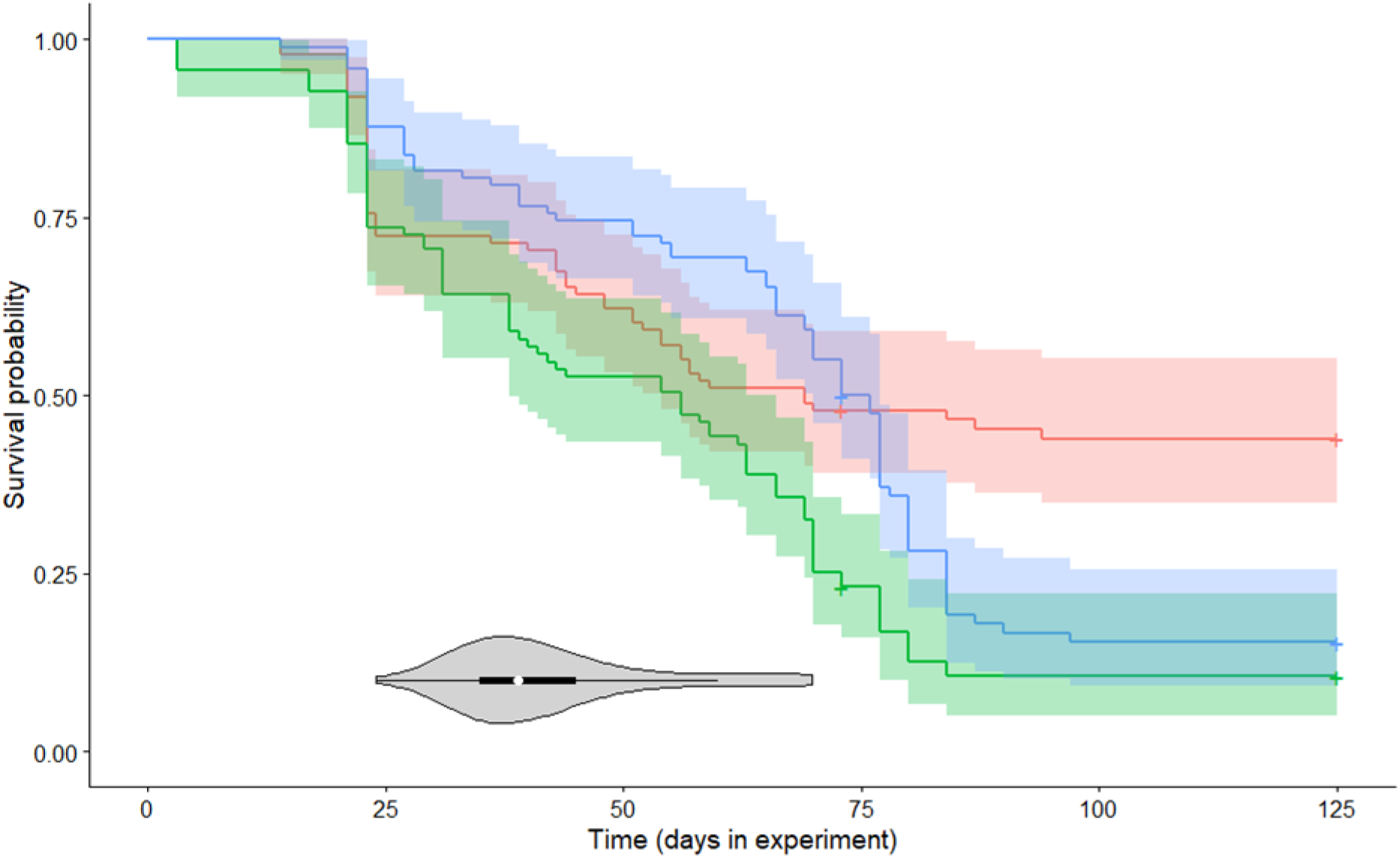
Survival over the experiment in the three treatments: control (red), Hungarian Bd isolate (green), Spanish Bd isolate (blue), with 95% confidence intervals. The violin plot shows the starting dates of metamorphosis (white dot: median, black box: interquartile range, grey curved areas: Kernel density plots). Ujszegi et al. Fig 1

**Table 2:**
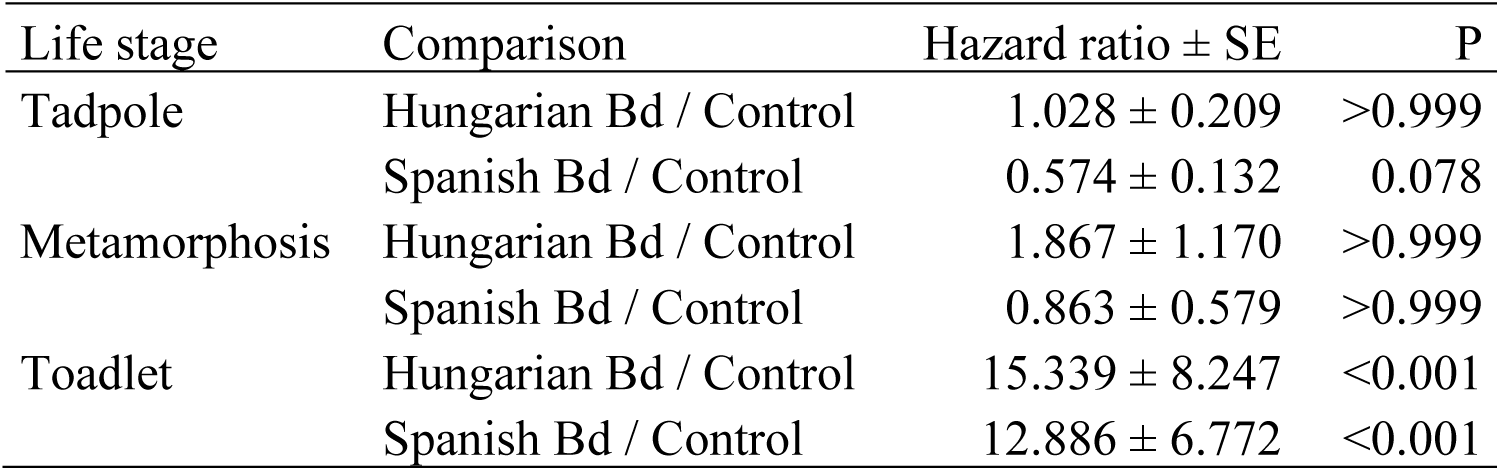
Effects of Bd treatments on mortality in three life stages, estimated from model 1.

| Life stage | Comparison | Hazard ratio ± SE | P |
| --- | --- | --- | --- |
| Tadpole | Hungarian Bd / Control | 1.028 ± 0.209 | >0.999 |
| | Spanish Bd / Control | 0.574 ± 0.132 | 0.078 |
| Metamorphosis | Hungarian Bd / Control | 1.867 ± 1.170 | >0.999 |
| | Spanish Bd / Control | 0.863 ± 0.579 | >0.999 |
| Toadlet | Hungarian Bd / Control | 15.339 ± 8.247 | <0.001 |
| | Spanish Bd / Control | 12.886 ± 6.772 | <0.001 |

Treatment with the Spanish Bd isolate significantly increased the frequency of sex reversal by genetic females into phenotypic males (Fisher’s exact test, *P* = 0.045, Table 3); we did not observe female-to-male sex reversal in the other two treatment groups. One of the three sex-reversed individuals in the Spanish Bd treatment group was the only surviving toadlet that had a detectable Bd load at the end of the experiment (Table 1). Additionally, a single toadlet in the control group showed male-to-female sex reversal (Table 3).

**Table 3:** Distribution of combinations of genetic and phenotypic sex across the treatment groups.

| Sex | Treatment group |  |  |
| --- | --- | --- | --- |
|  | Control | Spanish Bd | Hungarian Bd |
| Genetic females |  |  |  |
| Phenotypic females | 17 | 6 | 3 |
| Phenotypic males | 0 | 3* | 0 |
| Genetic males |  |  |  |
| Phenotypic males | 15 | 3 | 2 |
| Phenotypic females | 1† | 0 | 0 |

\*female-to-male sex reversal
†male-to-female sex reversal

## Discussion

Our data on Bd prevalence and infection intensity measured on individuals that died before the end of the study prove that our experimental exposure successfully infected larval toads. Interestingly, except for one individual with a very weak Bd load, juvenile toads that survived until the end of the experiment showed no detectable Bd load when dissected. Whether these individuals did not become infected during larval development, or cleared infection after metamorphosis remains unknown. Successful infection also manifested in higher mortality after metamorphosis of Bd-exposed individuals compared to the control group. This agrees with previous findings that the survival of common toads after metamorphosis can sharply decrease upon infection with Bd (Garner et al. 2009; Bielby et al. 2015); however, in our earlier experiments, we did not experience significant effect of Bd exposure with the same isolates on mortality of newly metamorphosed individuals originating from Hungarian common toad populations (Ujszegi et al. 2021; Kásler et al. 2022). This discrepancy may be explained by differences in the timing and/or duration of Bd exposure and the age of animals at Bd sampling. Furthermore, in the year of the present study, climatic conditions were extremely unfavourable for amphibian breeding in the spring, due to draught and short warm periods followed by repeated episodes of harsh frost. Consequently, the wild-caught tadpoles we used in the experiment had also been exposed to severe cold stress during their embryonic development. This probably contributed to the slow development and relatively high mortality in the control group (Beattie et al. 1991; Wersebe et al. 2019), and additionally to reduced Bd-tolerance, since high temperature variability can increase amphibian susceptibility to Bd infection (Raffel et al. 2013). In line with these considerations that mortality in the present study was likely due to multiple stressors, we found that the effect of Bd treatment on survival was time-dependent. In the early weeks of the experiment, when we suspect that mortality was mostly due to embryonic cold stress and perhaps the control broth we had to add into the tadpoles’ rearing water, survival did not differ significantly across treatment groups. In contrast, after metamorphosis, animals that had been exposed to Bd during larval development showed significantly higher mortality.

We found no effect of genetic sex on survival, although mortality was relatively high throughout the experiment and, therefore, would have allowed sex-specific differences to manifest. Furthermore, the effect of Bd treatments did not differ significantly between males and females. These results suggest that early-life mortality is not sex-dependent in common toads, similarly to other species (Bókony et al. 2021), and that the lethal effects of Bd infection are unlikely to skew sex ratios of young animals through sex-biased mortality. However, sex differences in susceptibility to infections and mortality risk can vary across age groups (Klein & Flanagan 2016), so it remains important to include both sexes and to investigate and report sex-specific responses in studies of Bd-effects in various life stages of amphibians.

Despite the facts that sex reversal can be caused by many types of stressors, and infections with pathogens and parasites are among the most important biological stressors, our study is the first to test the effect of pathogen exposure on sex reversal in any ectothermic vertebrate. Although the unexpectedly high mortality in our experiment constrains our sample size for investigating sex reversal, several lines of evidence suggest that Bd exposure may have interfered with sex determination and/or gonad development in our study animals. First, we found female-to-male sex reversal only in Bd-exposed animals. The stressors known to trigger sex reversal almost always do so in this direction in amphibians (Nemesházi & Bókony 2022), similarly to fish where a physiological link between endocrine stress and female-to-male sex reversal has been demonstrated (Castañeda-Cortés & Fernandino 2021). Second, out of the three female-to-male sex-reversed toadlets, one had detectable Bd load while all other toadlets were Bd-negative. Third, the proportion of genetic females that developed male phenotype (3 out of 9 toadlets in the Spanish Bd group) was remarkably high compared to the rare incidence of female-to-male sex reversal observed in common toads in earlier studies. In free-living populations, Nemesházi and colleagues (2022) found only one female-to-male sex reversal out of 349 adults in an earlier study, whereas no sex-ratio skew occurred in experiments that exposed large numbers of tadpoles to a heat wave (Ujszegi et al. 2022) or chemical contaminants (Bókony et al. 2020) under very similar laboratory conditions. These earlier results suggest that the common toad is relatively resistant against female-to-male sex reversal (Nemesházi & Bókony 2022). Therefore, despite the low sample size, our present finding that one-third of individuals exposed to the Spanish Bd isolate underwent female-to-male sex reversal is noteworthy, suggesting that Bd infection can induce sex reversal in juvenile toads. Interestingly, we did not experience this effect with the Hungarian Bd isolate, which might be because the sample size of surviving toadlets was even smaller in this group than in those exposed to the Spanish isolate. Alternatively, toad populations in Hungary may have adapted to the effects of the Bd strain present locally, including resistance against its potentially sex-reversing effect, but this speculation needs further investigation (e.g. see Nemesházi et al. 2022) for a study on resistance against chemically induced sex reversal). Additionally, we detected male-to-female sex reversal in one toadlet from the control group, although we cannot exclude the possibility of contamination in this case because we had no extra tissue sample in store from this individual to repeat genetic sexing.

In conclusion, our results indicate that Bd infection is unlikely to disturb the sex ratio of common toad populations through sex-specific mortality of infected tadpoles and young juveniles, but it may cause some genetic females to develop into phenotypic males. Even though the latter finding needs corroboration because of the relatively small sample size, it highlights that the relationship between sex reversal and Bd exposure deserves further attention in amphibians. Since the identification of sex reversal hinges on the availability of genetic sexing methods, which is currently limited to a handful of amphibian species (Nemesházi & Bókony 2022), a sex-reversing effect of Bd infection might be hidden in nature and could pose a serious threat for the already declining populations of several amphibian species.

## Acknowledgements

We are thankful to Dávid Herczeg, Dóra Holly, Szabolcs Hócza, Csenge Kalina, Anna Kraxner and Márk Szederkényi for their help during the experiment and data archiving, and Beata Rozenblut-Kościsty for help with interpreting histological images. We thank Éva Szita, Márk Z. Németh and all colleagues at the NÖVI Department of Plant Pathology for providing us with their microscopes and other lab facilities. Zoospore genomic equivalent standards were kindly offered by Judit Vörös and Jaime Bosch. The study was funded by the Lendület Programme of the Hungarian Academy of Sciences (MTA, LP2012-24/2012), and the National Research, Development and Innovation Office of Hungary (NKFIH, grants 135016 to V.B. and 124375 to A.H.). The authors were supported by the János Bolyai Research Scholarship of the Hungarian Academy of Sciences (to V.B. & A.H.), the New National Excellence Program of the Ministry for Innovation and Technology from the source of the National Research, Development and Innovation Fund (ÚNKP-21-5 and ÚNKP-22-5 to V.B. & A.H., ÚNKP-21-3 to A.K., ÚNKP-22-3 to E.B. and A.K., and ÚNKP-22-4 to J.U.), and the University of Veterinary Medicine Budapest (E.B. and V.B.). Experimental procedures were approved by the Ethical Commission of the Plant Protection Institute, and permissions were issued by the Government Agency of Pest County (PE-06/KTF/00754-8/2022, PE-06/KTF/00754-9/2022, PE-06/KTF/00754-10/2022, PE/EA/295-7/2018). The experiments were carried out according to recommendations of the EC Directive 86/609/EEC for animal experiments (http://europa.eu.int/scadplus/leg/en/s23000.htm). The authors have no conflict of interest to declare.

